# Externally driven attention to internal working memory content

**DOI:** 10.64898/2026.04.08.717206

**Authors:** Anna M. van Harmelen, Laurie Rol, Freek van Ede

## Abstract

To understand human thought and behaviour, it is vital to understand how internal representations ‘in mind’ interface with external sensations in the outside world. It is well-established that representations in working memory can involuntarily draw attention towards objects in the outside world with corresponding visual features. Here, we establish that this interface involves a two-way stream by demonstrating that external sensations also involuntarily draw attention to corresponding internal representations in working memory. We demonstrate this across four dedicated experiments in which an unpredictive external stimulus selectively colour-matched working-memory contents, while being completely irrelevant to the working-memory task. We provide converging evidence for this “externally driven internal attention” from tracking attentional dynamics over time through spatial biases in microsaccades as well as from memory performance. We further highlight that engagement with the external stimulus critically shapes the strength of this attentional capture from perception to memory. Together, our findings delineate the properties and consequences of this underexplored attentional pathway, opening novel avenues for research at the interface of perception and memory, and their disorders.

## Introduction

Selective attention is the essential process by which the brain selects and prioritises information. Historically, research on selective attention has concentrated on external attention^1–6^, where the *target* of attention concerns sensory information from the external environment. Within this long-standing line of research, a foundational distinction is made with respect to the *source* of attention, which can be endogenous (voluntary or goal-driven) or exogenous (involuntary or stimulus-driven). Besides selective attention targeting external sensations, our attentional focus can also be directed internally to the contents of working memory^7–13^.

Unlike the study of external selective attention, research on internal selective attention has predominantly been studied by prompting the prioritisation of working-memory contents in a voluntary, goal-directed manner^14^. However, when differentiating the *source* (endogenous/exogenous) and the *target* (internal/external) of selective attention, as we illustrate in **Figure 1**, it becomes conceivable that internal selective attention may also be subject to exogenous (involuntary or stimulus-driven) influences. Specifically, external stimuli may involuntarily draw attention to internal representations in working memory that share one or more of their features. In this article we will refer to this as “externally driven internal attention”. A small but growing body of literature has recently started to provide initial evidence for this form of selective attention^15–22^. Yet, compared to the other three categories of attention illustrated in **Figure 1**, these exogenous influences on attention within working memory remain distinctly underexplored.

**Figure 1:**
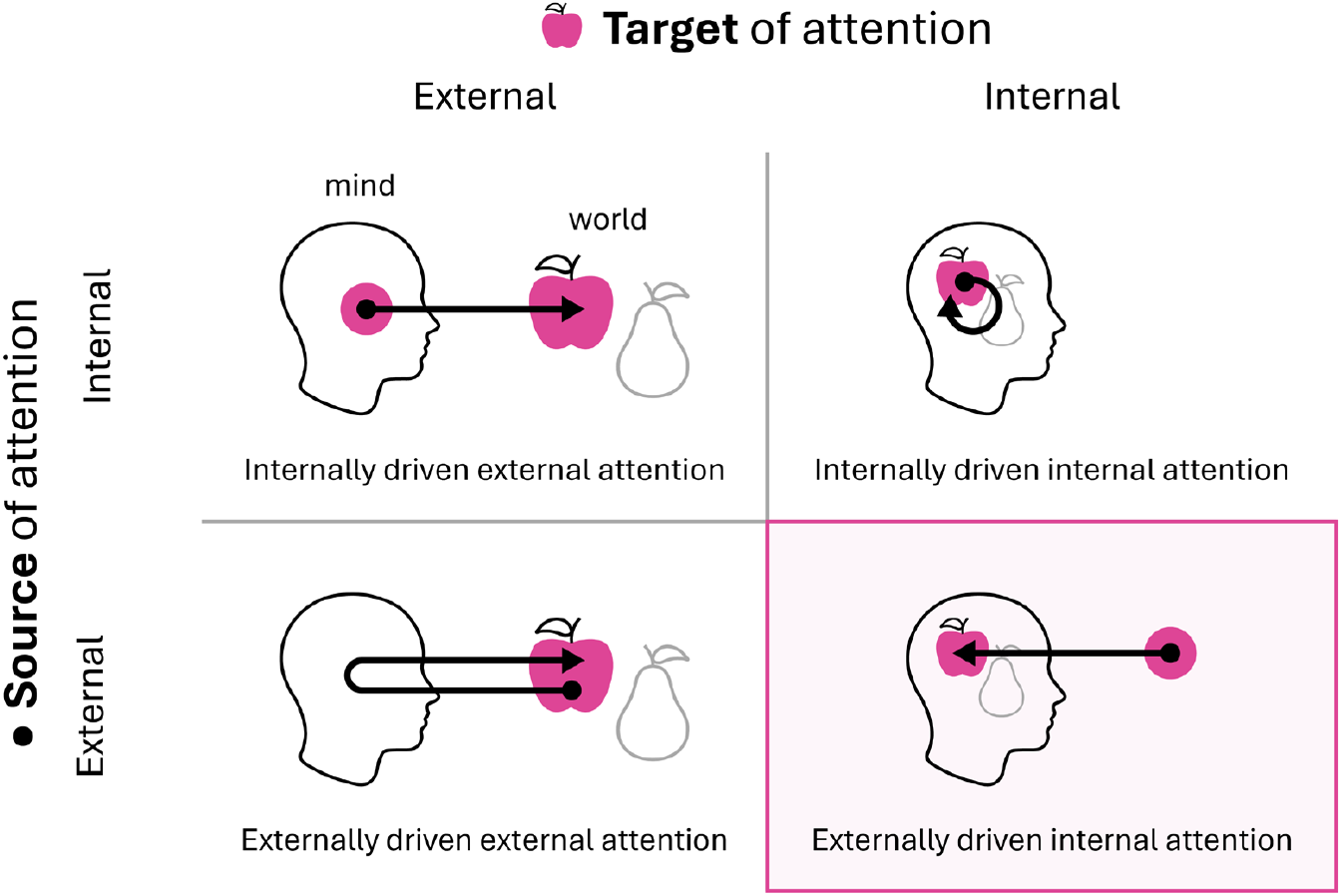
taxonomy of attention based on its source and target. The taxonomy shows that both the source and target of attention can be either external or internal. Selective attention can be deployed in a voluntarily goal-driven manner (internal source) or involuntarily stimulus-driven manner (external source), and can steer selection and prioritisation among external sensations (external target) or internal representations (internal target). This article focuses on the comparatively underexplored form of attention whereby an external stimulus involuntarily drives internal selective attention among working-memory contents (highlighted in pink).

Two main factors may account for the comparative underrepresentation of externally driven internal attention in the literature. First, it is possible that the topic has simply been largely overlooked. Compared to research on external selective attention, the study of internal selective attention has emerged relatively recently^13,14^, and may therefore contain underexplored areas. A second possibility is that exogenous influences on internal selective attention within working memory have in fact been amply investigated, but that the observed effects were weak or inconsistent. This would result in relatively few published findings, considering the well-known publication bias or ‘file-drawer problem’^23–25^. Indeed, the available studies report relatively modest effects^15–22^.

We propose that the relatively modest effects and scarcity of published studies on externally driven internal attention may stem from a dependence on specific experimental parameters. It is useful here to draw a parallel with complementary literature on memory-guided external attention: this field has examined how items held in working memory involuntarily draw attention to matching external stimuli^26–32^. Crucially, this effect is typically studied in the context of an intervening task, such as a visual search task in which the external stimuli cannot be ignored. By contrast, the aforementioned studies investigating externally driven internal attention have typically employed external stimuli that were entirely task-irrelevant and therefore, in principle, ignorable. Indeed, evidence shows that participants can effectively suppress distracting external information during working-memory maintenance^33–35^. We theorise that the absence of any mandatory engagement with the external stimuli in prior studies may have considerably constrained their capacity to influence internal selective attention.

In the current study, we provide clear evidence of an exogenously driven allocation of internal selective attention amongst the contents of visual working memory. We show that internal attention can be strongly driven by memory-matching external stimuli that are actively processed, yet completely irrelevant for the working-memory task. We demonstrate the stimulus-locked nature of this influence through spatial biases in microsaccades in the direction of matching internal visual representations, as held within the spatial layout of working memory. We further show that this underexplored form of selective attention is constituted by both benefits to matching and costs to non-matching working-memory contents. Finally, we substantiate our hypothesis that engagement with the external stimulus is a key factor in drawing attention involuntarily to matching working-memory contents.

## Results

Healthy human volunteers performed a task designed to address whether and how an external stimulus may involuntarily draw attention to a feature-matching internal representation held in working memory (**Fig. 2A**). Participants held two visual items (tilted bars) in working memory until a coloured response dial was shown on screen, prompting participants to reproduce the orientation of the colour-matching item in working memory.

**Figure 2.**
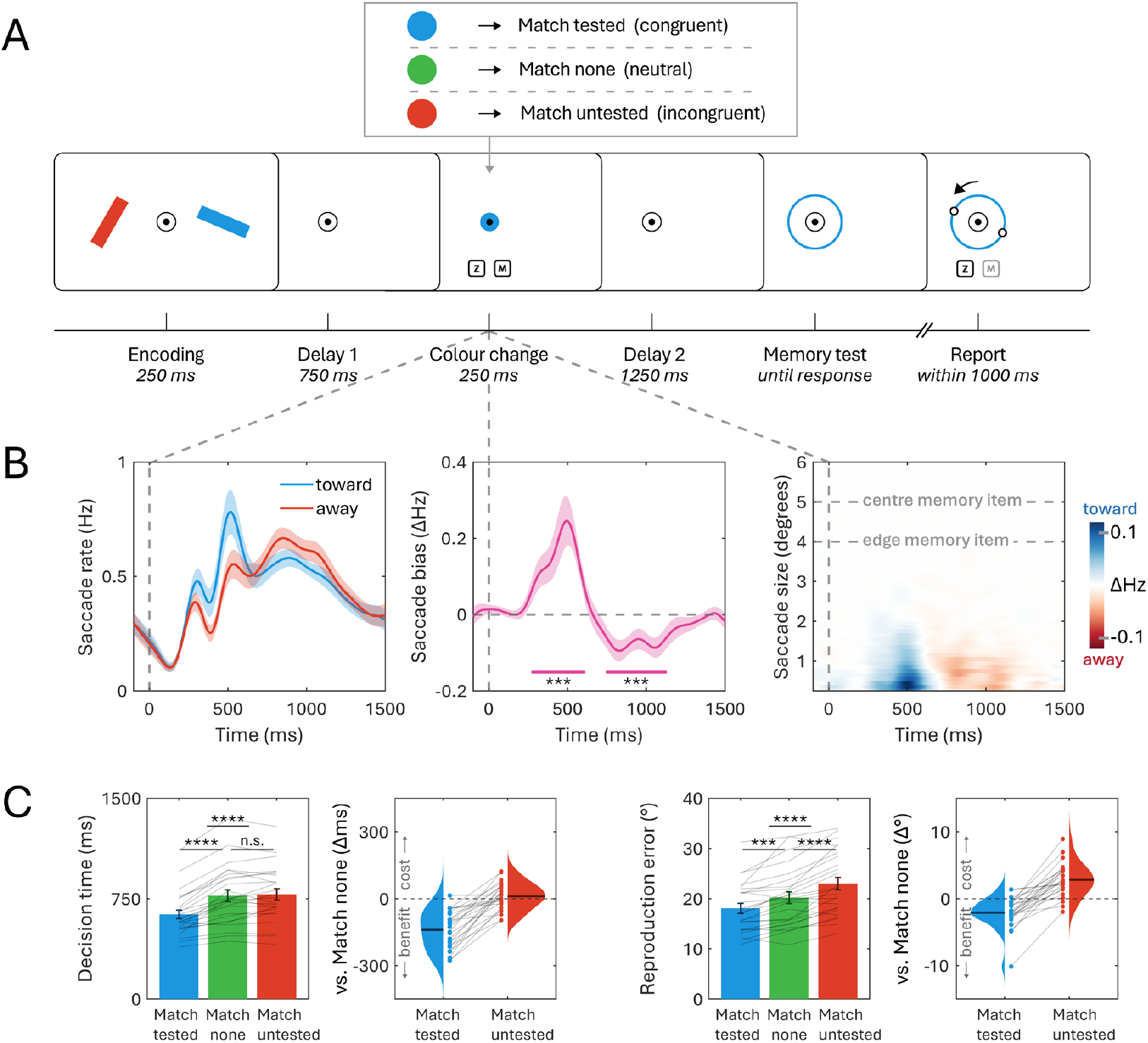
An external colour stimulus can involuntarily draw internal attention to matching working-memory content. (A) Schematic of the working-memory task designed to address whether and how an external stimulus may involuntarily draw attention to a feature-matching internal representation held in working memory. This colour change was completely uninformative as to which item would be tested at the end of a trial. Participants were required to evaluate whether the colour change matched a predetermined colour term and responded with a button press (buttons are shown here for illustration purposes and were not shown on screen during the experiment). Colours shown serve only as an example, for the experimental colours, see the methods section. (B) From left to right: time course of saccade rates toward and away from the colour-matching memory item; time course of the saccade bias reflecting internal selection as measured by saccade rate (toward – away); time course of the saccade bias as a function of saccade size. For reference two dashed horizontal lines are shown to indicate the original location of the centre and closest border of the memory item. (C) Behavioural performance (left: decision times, right: reproduction errors) as a function of whether the colour change matched the eventual tested memory item, the untested memory item, or was neutral (matched neither memory item). Bars reflect the mean behavioural performance, with whiskers indicating the standard error of the mean. Grey lines indicate individual participants’ data. Violin plots show the match-tested and match-untested conditions compared to the neutral condition. Within the violin plots, dark lines show the mean difference and circular markers indicate individual participants. Throughout the entire figure, the following significance levels were used: *: p< 0.05, **: p < 0.01, ***: p < 0.001, ****: p < 0.0001.

To study the involuntary influence of an external stimulus on internal attentional allocation in working memory, the central fixation marker briefly changed colour during the delay period. Throughout the entire experiment, this colour change was entirely unpredictive of which item would be tested during the working-memory task. Importantly, to ensure participants engaged with the uninformative colour change and could not simply ignore it, participants were required to perform a perceptual evaluation of this colour change. This intervening task was entirely irrelevant to the working-memory reproduction task. Participants were asked to simply perform a double-button press when the colour change matched a predetermined colour, as specified by a colour term at the beginning of the block (see the methods for details). We return to the effect of this manipulation later in the results section.

We conducted multiple complementary experiments that we present in succession. We first show that an unpredictive external stimulus can involuntarily draw attention to matching working-memory content. Next, we replicate these effects while ruling out a memory-based verification strategy for the intervening task, thereby corroborating the involuntary nature of the effects. Finally, we show that engagement with the unpredictive external stimulus is a key factor in shaping the strength of the involuntary, stimulus-driven, influence of an external stimulus on internal attentional selection among the contents of working memory.

### An external colour stimulus can involuntarily draw internal attention to matching working-memory content

To track internal attentional allocation in response to the external colour change during the working-memory delay interval, we employed a recently developed peripheral marker of internal attentional shifts. Previous work has reliably established spatial biases in microsaccades as a marker of attention shifts among visual contents held within the spatial layout of working memory^36–39^. Here, we leveraged the microsaccade bias in response to the colour change to examine whether this colour change prompted an involuntary attention shift to the colour-matching item held in working memory.

**Figure 2B (left panel)** shows the time-resolved saccade rates following the unpredictive colour change separately for saccades directed toward or away from the original location of the colour-matching memory item. Note that the colour change and the upcoming working-memory test were always presented centrally. Consequently, any spatial bias reported here implies an attentional shift among the contents of working memory. We found a robust bias in saccade direction towards the original location of the colour-matching item, as quantified by the difference between toward and away saccades (**Fig. 2B, middle panel**: cluster p=0.0003). This initial attraction bias was followed by a bias in the opposite direction (cluster p=0.0002), likely reflecting corrective saccades aimed to return to central fixation. Finally, we analysed the sizes of the saccades contributing to this saccade bias, which confirmed the effect was predominantly driven by saccades in the “micro”-range (**Fig. 2B, right panel**), consistent with earlier work^36,39^. Together, these data show a stimulus-driven biasing of microsaccades within the spatial layout of visual working memory, signifying that the colour change involuntarily draws attention to matching working-memory content.

Complementing this time-resolved marker of internal attentional shifting, we could also corroborate that our unpredictive stimulus had consequences for performance on the working-memory reproduction task. We accordingly analysed decision times and reproduction errors on the reproduction task as a function of whether the unpredictive colour stimulus matched the item that was eventually tested, or the other item. As shown in **Figure 2C**, responses on the working-memory reproduction task were faster (main effect of condition: p<0.0001; match tested vs. untested: t(24)=8.64, p_Bonferroni_<0.0001, d=0.78) and more accurate (main effect of condition: p<0.0001; match tested vs. untested: t(24)=9.06, p_Bonferroni_<0.0001, d=0.84) when the colour change matched the eventual tested item instead of the other item. This provides further evidence that attention had been allocated to the colour-matching memory item.

We also confirmed whether participants performed the intervening task sufficiently, which was introduced to ensure that participants could not simply ignore the unpredictive colour stimulus. Participants performed this intervening task well, with an average hit rate of 91.24±10.31% (M±SD), and an average correct-rejection rate of 95.60±3.39% across participants.

### Externally driven internal attention yields benefits to matching and costs to non-matching working-memory content

The performance data from trials in which the colour change did not match either memory item enabled us to assess whether externally driven internal attention reflects benefits to matching items and/or costs to other items held in working memory. These trials served as a neutral baseline when assessing potential costs and benefits from the behavioural performance data. **Figure 2C** shows that both decision times and reproduction errors were significantly reduced in the match-tested condition compared to the neutral condition (decision time: t(24)=8.30, p_Bonferroni_<0.0001, d=0.71; reproduction error: t(24)=4.53, p_Bonferroni_=0.0004, d=0.37), confirming a benefit to colour-matching items. In contrast, decision times were not significantly affected when the colour change matched the untested item (t(24)=0.86, p_Bonferroni_=1.00, d=0.05, BF_01_=3.41), but errors did increase in comparison to the neutral condition (t(24)=5.64, p_Bonferroni_<0.0001, d=0.45). Together, these findings show that externally driven internal attention can induce both a benefit to the representation of the matching memory item, and a cost to the other memory item.

### Replication of findings while ruling out a memory-based verification strategy

In the intervening task of Experiment 1, participants judged whether a central colour change matched a predetermined colour (for example green, as in **Fig. 3A**). This predetermined colour always corresponded to the neutral colour not used for either memory item. As illustrated in **Figure 3A**, this design creates the possibility that participants could have, in principle, solved the intervening task via a memory-based verification strategy: rather than evaluating the colour change directly, they could deduce the correct response by checking whether the colour matched either or neither memory item. Such a strategy of revisiting working memory in service of the intervening task, though seemingly cumbersome, could potentially explain the observed microsaccade bias and accompanying performance effects. We therefore conducted a follow-up experiment that was identical to Experiment 1, except for the inclusion of an additional neutral colour. As shown in **Figure 3A**, under this design, a memory-based verification strategy alone no longer provides sufficient information to perform the intervening task, since participants were instructed to respond only to a specific neutral colour, rather than any neutral colour. Experiment 2 therefore enabled us to ensure that the original effects replicate, while also ruling out a memory-based verification strategy as an alternative explanation.

**Figure 3.**
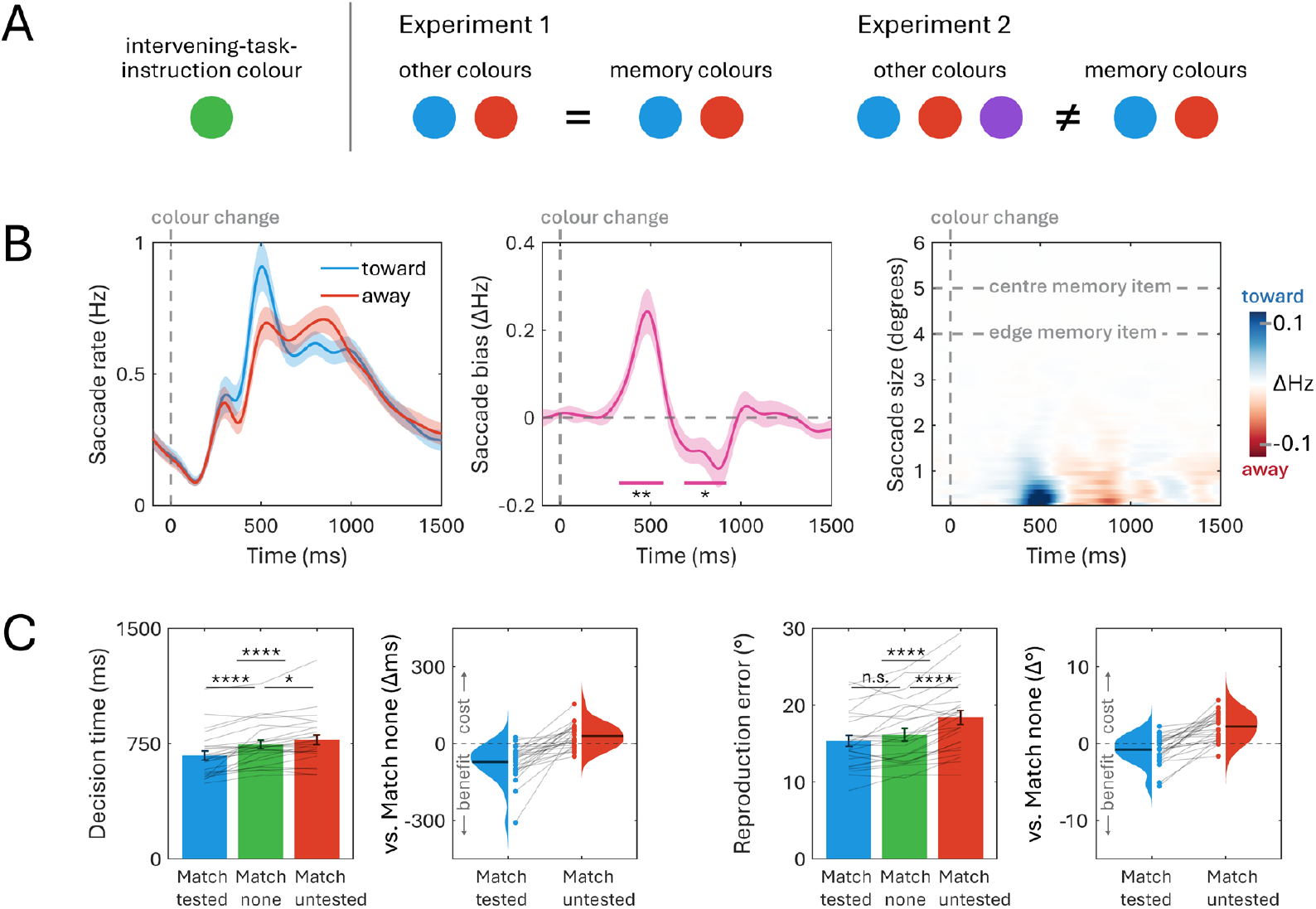
Replication of findings while ruling out a memory-based verification strategy. (A) Illustration of how the design of Experiment 1 creates the possibility that participants could have, in principle, solved the intervening colour-evaluation task through a memory-based verification strategy: rather than evaluating the colour change directly, they could deduce the correct response by checking whether the colour matched either or neither memory item. By adding a second neutral colour, this was no longer viable in Experiment 2. Colours shown serve only as an example, for the experimental colours, see the methods section. (B) From left to right: time course of saccade rates toward and away from the colour-matching item; time course of the saccade bias reflecting internal selection as measured by saccade rate (toward – away); time course of the saccade bias as a function of saccade size. For reference two dashed horizontal lines are shown to indicate the original location of the centre and closest border of the memory item. (C) Behavioural performance (left: decision times, right: reproduction errors) as a function of whether the colour change matched the eventual tested memory item, the untested memory item, or was neutral (matched neither memory item). Bars reflect the mean behavioural performance, with whiskers indicating the standard error of the mean. Grey lines indicate individual participants’ data. Violin plots show the match-tested and match-untested conditions compared to the neutral condition. Within the violin plots, dark lines show the mean difference and circular markers indicate individual participants. Throughout the entire figure, the following significance levels were used: *: p< 0.05, **: p < 0.01, ***: p < 0.001, ****: p < 0.0001.

We first verified that participants were still able to perform the intervening task with the additional neutral colour. As in the previous experiment, participants were able to do the task well, with an average hit rate of 95.04± 4.32% (M±SD) and an average correct-rejection rate of 96.36±3.50% across participants. In fact, there was no significant difference in performance on this intervening task between Experiment 1 and Experiment 2 (hit rate: t(24)=1.70, p=0.096, d=0.47, BF_01_=1.10; correct-rejection rate: t(24)=0.77, p=0.444, d=0.22, BF_01_=2.77).

**Figure 3** shows that the results from Experiment 1 were closely replicated in Experiment 2: we again observed a robust microsaccade bias towards the original location of the colour-matching item (**Fig. 3B, middle panel**: cluster p=0.002), followed by a return bias in the opposite direction (cluster p=0.015). This was again driven predominantly by saccades below 1 degree visual angle (**Fig. 3B, right panel**). Similarly, responses on the working-memory task were again faster (main effect of condition: p<0.0001; match tested vs. untested: t(24)=5.96, p_Bonferroni_<0.0001, d=0.65) and more accurate (main effect of condition: p<0.0001; match tested vs. untested: t(24)=6.91, p_Bonferroni_<0.0001, d=0.72) when the colour change matched the eventual tested item compared to the other item. Furthermore, a significant benefit was again found for decision time in the match-tested condition compared to the neutral conditions (t(24)= 5.16, p_Bonferroni_<0.0001, d=0.49), but no significant benefit was found for reproduction error in the match-tested condition compared to the neutral conditions (t(24)= 2.12, p_Bonferroni_=0.134, d=0.20). We did observe significant costs for the match-untested condition compared to the neutral conditions, for both decision times (t(24)= 3.24, p_Bonferroni_=0.011, d=0.20) and reproduction errors (t(24)= 6.28, p_Bonferroni_<0.0001, d=0.50). Overall, these results confirm that externally driven internal attention can lead to both benefits to matching and costs to other memory items.

Experiment 2 thus replicated the main findings from Experiment 1, confirming that an unpredictive colour stimulus can draw attention to matching working-memory content. Experiment 2 additionally confirmed that this effect persists when ruling out a memory-based verification strategy as an alternative explanation, which corroborates the involuntary nature of the reported effects.

### Stimulus engagement critically shapes externally driven internal attention

Lastly, we investigated the importance of engaging with the colour change for involuntarily drawing attention to the colour-matching item in working memory. To do so, we compared the combined data from Experiment 1 and Experiment 2 to two previously collected datasets with an identical experimental set-up, with the only exception that the colour change did not require any perceptual evaluation or response, and could therefore simply be ignored (as has been the default in prior studies in this line of research^15–21^). Note that in both cases the colour change was equally unpredictive and equally irrelevant to the primary working-memory reproduction task.

**Figure 4A** shows that the spatial microsaccade bias was considerably larger when participants were required to engage with the external colour change than when it could be ignored. When engagement was required (data collapsed across Experiments 1 and 2), significant microsaccade biases were observed both toward (**Fig. 4A**: cluster p<0.0001) and away (cluster p= 0.002) from the original location of the colour-matching item, consistent with the results reported above. When the colour change could be disregarded, the microsaccade bias was less clear, but could still be observed (**Fig. 4A**: cluster p=0.002). Critically, when directly comparing the engage (Experiments 1-2) and disregard (Experiments 3-4) conditions, the microsaccade bias was significantly larger when it was required to engage with the colour change (cluster p=0.008).

**Figure 4.**
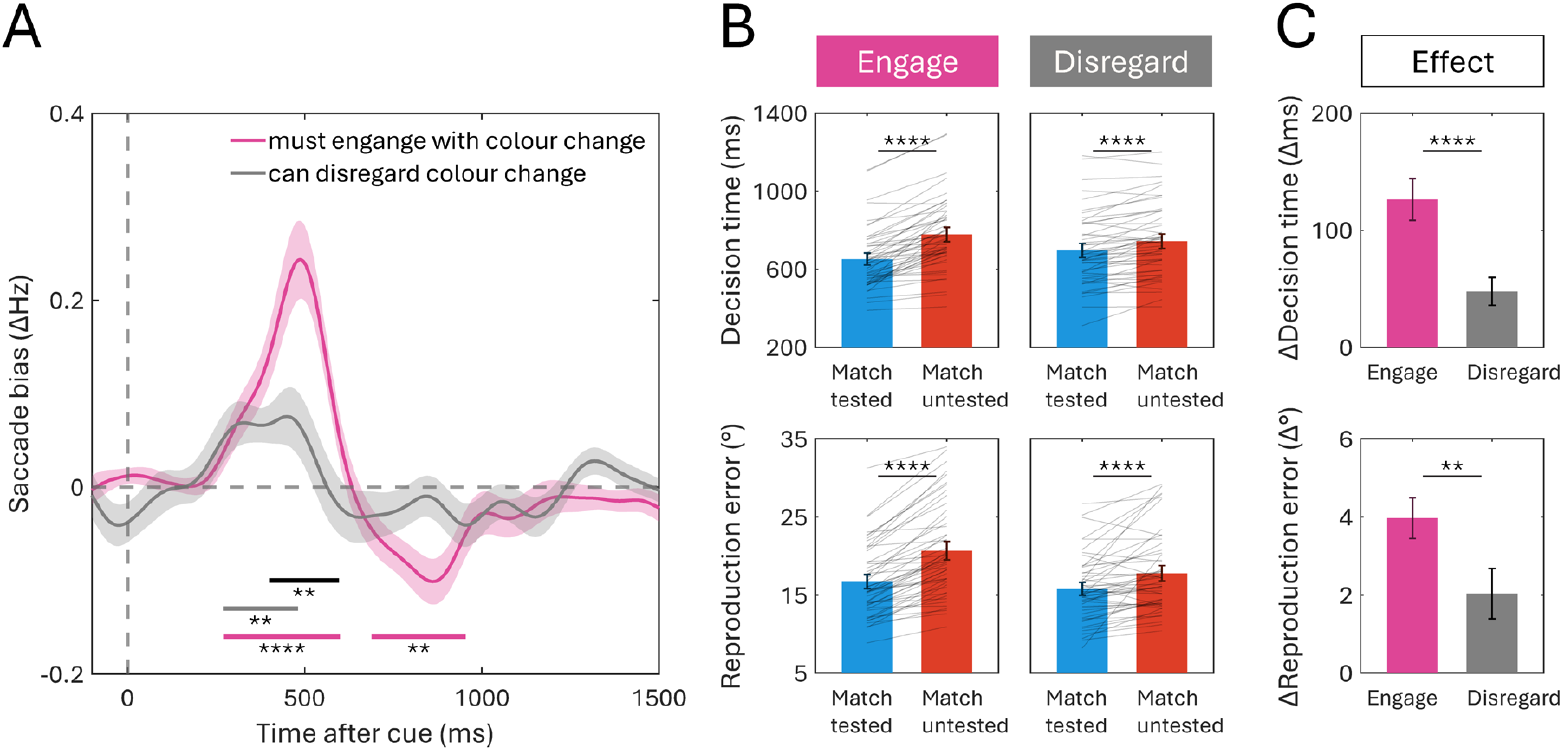
Stimulus engagement critically shapes externally driven internal attention. (A) Time courses of saccade biases reflecting internal selection as measured by saccade rate (toward – away) when engagement with the colour change was required (pink) and when the colour change could be disregarded (grey). Horizontal lines mark the timepoints with a significant difference against zero (pink and grey), or between the two shown conditions (black). (B) Behavioural performance (decision times above, reproduction errors below) when engagement with the colour change was required (‘Engage’) and when the colour change could be disregarded (‘Disregard’). Bars reflect the mean behavioural performance, with whiskers indicating the standard error of the mean. Grey lines indicate individual participants’ data. (C) The comparison between the internal-attention effects (match tested vs. match untested) in the ‘engage’ and ‘disregard’ conditions. Bars reflect the mean effect, with whiskers indicating the standard error of the mean. Throughout the entire figure, the following significance levels were used: *: p< 0.05, **: p < 0.01, ***: p < 0.001, ****: p < 0.0001.

The performance data mirrored these microsaccade findings. **Figure 4B** shows the performance effects for the corresponding conditions, quantified again as the difference between match-tested and match-untested items. These effects were significant both when the colour change had to be evaluated and when it could be disregarded, but they were significantly larger in the former case. Consistent again with the results described above, significant performance differences were observed when engagement was required, for both decision times (**Fig. 4B**: t(49)=10.04, p<0.0001, d=0.74) and reproduction errors (**Fig. 4B**: t(49)=10.74, p<0.0001, d=0.74). Performance effects were smaller when the colour change could be disregarded, but still significant, for both decision times (**Fig. 4B**: t(49)=5.68, p<0.0001, d=0.26) and reproduction errors (**Fig. 4B**: t(49)=4.47, p<0.0001, d=0.44). Critically, performance effects on the working-memory task were significantly larger when engagement with the colour change was required by the intervening task than when it could be disregarded (**Fig. 4C**), for both decision times (t(98)=5.19, p<0.0001, d=1.03) and reproduction errors (t(98)=3.30, p=0.001, d=0.65).

Together, the microsaccade and performance results reveal that stimulus engagement is a critical variable in shaping the strength and robustness of exogenously driven internal attention shifts among the contents of working memory.

## Discussion

Our results demonstrate that an unpredictive external stimulus can involuntarily draw attention to matching internal contents in working memory. This is supported by converging evidence from stimulus-locked biases in microsaccades towards matching working-memory content and from modulations of performance on the ensuing working-memory task. We further show that external stimuli can cause both benefits to matching working-memory content and costs to other working-memory content. Finally, we demonstrate that the strength of externally driven internal attention hinges on engagement with the external stimulus that initiates the attentional shift.

The considerable effect of engagement raises an important question about the degree to which externally driven internal attention is automatic. In the present study, we uniquely studied externally driven internal attention while participants were required to actively engage with a colour change that was irrelevant to the working-memory content and the upcoming working-memory task. For this reason, we characterise the effect as involuntary, since attention being drawn to the colour-matching memory items provided no strategic advantage. At the same time, the observation that engagement with the unpredictive external colour stimulus modulated the effects on both microsaccades and performance suggests that externally driven internal attention, while involuntary, is not triggered in a fully automatic fashion. We therefore provide evidence for a form of *contingent* automaticity in externally driven internal attention, paralleling related findings and interpretations from the complementary literature on the involuntary memory-guided allocation of external attention^26–32^.

Why might external stimuli automatically draw attention to matching internal working-memory representations? A useful parallel can again be drawn with the literature on memory-guided externally directed attention, which has argued that “involuntary” interactions from working memory to perceptual attention may reflect mechanisms that are ordinarily adaptive in everyday behaviour ^31^. For example, in daily life, visual contents of working memory often correspond to what is actively being searched for in the external world. Consequently, when an item is held in working memory, we may be naturally drawn to matching stimuli in the world to help us find them, even if this is not adaptive in a given laboratory task. A similar process may be at work in our present task. Under natural circumstances, when a possible target in the external world is detected, this may naturally invoke an internal verification process to determine whether the external stimulus is what we are looking for (see ^39^). In our task, the external stimulus was not the object of a visual search, but this verification process may nonetheless be triggered naturally. Although speculative, it is conceivable that the process we uncover here may be purposeful under natural circumstances. It might therefore be such a habitual response that it still occurs in contexts where it is no longer functional, as in our experimental task.

Our work complements and extends existing research on the same underexplored phenomenon of externally driven internal selective attention among the contents of working memory^15–22^. Adding to existing work, we provide three main advances. First and foremost, we establish that engagement with the external stimulus critically shapes the strength of the involuntary attentional allocation to matching working-memory content. Second, we demonstrate that the involuntary attentional allocation in working memory by an external stimulus occurs directly following stimulus onset, as shown by tracking attentional dynamics over time through spatial biases in microsaccades (for complementary neural evidence, see also ^22^). Third, we show that externally driven internal attention entails both benefits for matching working-memory content and costs for other memory content. In addition to these main advances, we also extend two closely related sets of studies in relevant ways. First, in a series of studies^20–22^, Fuentes-Guerra et al. reported effects of external stimuli that matched working-memory content in their spatial location. Here, we show such effects following a non-spatial colour stimulus. Second, earlier work from our lab^15^ employed colour cues as well, but additionally included blocks in which the external stimuli were informative and where participants therefore got used to using the external stimuli for the working-memory task. Here, we demonstrate that external stimuli can draw attention to colour-matching working-memory contents even when these external stimuli are completely unpredictive throughout the entire experiment. Taken together, our results thus contribute several novel conceptual insights into externally driven internal attention.

Our findings complement, but should also be distinguished from, several other related lines of prior work. First, in our paradigm the external stimuli (“cues”) were completely irrelevant to the working-memory task. This contrasts with multiple literatures in which internal representations are selected because they are more likely to become relevant (e.g., informative retrocues^8,13^), have higher value (e.g., learned reward associations^40,41^), are brought to mind based on explicit instructions (e.g., refreshing cues^42,43^), or are necessary to perform a secondary task (e.g., incidental cues^44,45^). In this respect, our findings more closely resemble “pinging” approaches that use external inputs that are, like ours, completely irrelevant to the working-memory task^46^. However, in contrast to conventional pinging studies, in our case the external stimulus selectively matched only one memory item. Second, our findings differ from the “retroperception effect”, in which cues are presented much earlier after encoding^47^ (see also ^48^). In our task, the external colour change occurred much later (750 ms after stimulus offset), suggesting that the external influence over internal attention we report occurs in working memory and not iconic memory. Third, building on this idea of memory reactivation, our results are conceptually related to targeted memory reactivation studies^49,50^ that typically focus on reactivation of long-term memory traces during sleep. In contrast, we studied selective “reactivation” of working-memory traces during wakefulness. Fourth, actions made to specific locations have been shown to also involuntarily draw attention to spatially congruent working-memory contents^51–53^. In our study, in contrast, internally directed selective attention was purely stimulus driven, and triggered in a non-spatial manner following central colour stimuli. Overall, our findings connect to ample prior research directions, while also complementing each in relevant ways.

Our findings also leave interesting avenues for future research. One envisioned line of future research could explore the neural mechanisms underlying the here-reported processes, noting that our microsaccade findings already suggest involvement of the oculomotor system (in line with ample prior studies linking the oculomotor system to covert and internal attention^54–56^. It would also be valuable to examine whether the current findings generalise to other cue types beyond colour, to different forms or levels of stimulus engagement, and to other memory systems. Lastly, it would be important to translate these findings to more ecologically valid experimental settings that better approximate daily life.

Taken together, across a series of complementary experiments, our findings firmly establish externally driven internal attention among working-memory content and provide an experimental approach to track it through biases in microsaccades. Furthermore, by unveiling how engagement with the external stimulus can considerably amplify this involuntary form of internal attention, our findings pave the way to delineate the properties and mechanisms of this distinctly underexplored attentional pathway – and how it may be altered across development, disorders, and disease.

## Methods

All experimental procedures were reviewed and approved by the ethics committee of the Vrije Universiteit Amsterdam. Each participant provided written informed consent prior to participation and either received participation credits or was reimbursed with 10 euros/hour or 12.50 euros/hour (after institutional policy changes).

### Participants

The primary experiments we report on here were two dedicated experiments that only differed in the number of neutral colours (detailed below under ‘*Experimental set-up*’). A sample size of twenty-five volunteers per experiment was set a-priori, based on previous publications from our lab with similar experimental designs and outcome measures (e.g., ^15,38^).

All participants were recruited from the Vrije Universiteit Amsterdam. Twenty-six healthy human volunteers participated in Experiment 1, one of which was excluded post-hoc because on 42% of trials they responded the maximum rotation of ± 90°, indicating that they did not understand that the response consisted of an initial forced choice in the correct direction. Their z-score on this behaviour (compared to the remaining participants) was 4.7. This resulted in the a-priori determined sample size of twenty-five participants (age range: 18-29; 21 women, 3 men and 1 non-binary; 22 right-handed). Twenty-eight healthy human volunteers participated in Experiment 2, two of which were excluded because they did not perform the full experiment, one of which was excluded because the eye-tracking data were deemed of too poor quality due to both a high noise level in the Y-channel and an excess of blinks. This resulted in the a-priori determined sample size of twenty-five participants (age range: 18-32; 20 women, 5 men; 18 right-handed). Recruitment for both experiments was performed independently, with the constraint that participants were not allowed to participate in both experiments.

To gain additional insight into our findings, we also compared the data from these two dedicated experiments to condition-specific data from two unpublished experiments that we had collected previously (referred to here as Experiments 3 and 4). We return to this under ‘*Comparison to comparable conditions in prior unpublished experiments*’ where we also list the relevant participant demographics of those experiments.

### Task and procedure

Experiment 1 and Experiment 2 employed an identical task, with one key difference that we return to later. The task consisted of a visual-working-memory task that required participants to remember two visual items and reproduce the orientation of one of them at the end of the working-memory delay. As shown in **Figure 2A**, two tilted bars with different colours were shown on either side of the central fixation marker for 250 ms, followed by a 2250 ms delay. The critical manipulation occurred after the first 750 ms of the delay, when the central fixation marker briefly changed colour for 250 ms. After this, the central fixation marker changed back to the original colour for the final 1250 ms of the delay.

The colour change of the central fixation marker was entirely unpredictive of which memory item would be tested at the end of a trial. However, to ensure participants would process this visual event, participant were instructed to perform an intervening task, which consisted of a perceptual evaluation of this colour change. This intervening task was entirely irrelevant to the working-memory reproduction task. At the start of each block, the intervening task instructions were presented, and could either be “respond when {colour}” or “respond when NOT {colour}”, where “{colour}” was the colour term for the neutral colour that was determined per participant and counterbalanced across participants (see ‘*Experimental set-up*’ for details). This colour was never used as the colour for either memory item for that participant. The intervening task response was a double-button press of both the ‘z’ and ‘m’ keys (to avoid having to switch response keys, we used the same keys that were also used to give the tilt response on the working-memory task, though for that participants always used either key, not both). For example: if a participant’s task-relevant neutral colour was green, and the block instructions were “respond when green”, then the correct response to a green central fixation marker would be to press both ‘z’ and ‘m’ keys concurrently. The correct response to a blue, orange or magenta central fixation marker would be to withhold a response. If the block instructions were “respond when NOT green”, then a green central fixation marker would not warrant a response, and a blue, orange, or magenta central fixation marker would require a response of both ‘z’ and ‘m’ keys.

After the delay period, a coloured circular response dial was shown on screen centred on the black-and-white central fixation marker, indicating that participants should now reproduce the orientation of the colour-matching item from working memory. Participants were instructed to make the response dial match up with the memorised orientation of the indicated item, as precisely as possible. The coloured response dial was shown until the participant finished their response. At response initiation (key press), an orientation-indicator appeared on the dial at a 0°, upright, angle, and kept turning until response termination (key release). The eventual position of the response dial was the responded orientation. Pressing ‘m’ made the response dial move clockwise, while pressing ‘z’ made the response dial move anti-clockwise. The response dial moved at a speed of 90°/s while the keyboard press was sustained, until a maximum of exactly 90°, after which the response was automatically terminated. It was not possible to release the response key to change the direction of the response. This forced participants to decide whether to report a clockwise or anti-clockwise orientation before pressing either key.

Both experiments contained 16 blocks of 48 trials each. 8 blocks were “respond when {colour}”-blocks, and 8 blocks were “respond when NOT {colour}”-blocks. For all analyses, we collapsed across these two block types. Before starting the experiment, participants first practiced using the response dial in isolation and then practiced entire trials from both block types until they felt acquainted with the task.

### Experimental set-up

Both tasks were programmed in Python 3.8.17 using the PsychoPy library^57^ to generate the stimuli. Participants were seated with their head in a chin-rest at approximately 70 cm in front of a 24-inch LCD monitor, with a 1920 x 1080 pixel resolution and a 239 Hz refresh rate.

For both Experiment 1 and Experiment 2, bar stimuli were presented on a dark grey background (rgb: 64, 64, 64). The bars were 0.6 x 4 degrees visual angle (width x height), presented at a 6 degree distance to the central fixation marker (centre to centre). The fixation marker was 0.35 degrees visual angle in diameter and near-white (rgb: 234, 234, 234), and contained a central fixation dot with a 0.1 degree diameter, which was black (rgb: 0, 0, 0). The orientations of the bar stimuli were randomly generated from a 160°-space, where orientations from -85 to -5 and 5 to 85 degrees were allowed, to avoid (near-)cardinal orientations.

Three colours were used for Experiment 1: blue (rgb: 19, 146, 206), green (rgb: 101, 148, 14), orange (rgb: 238,104, 60). All three colours were calibrated prior to the experiment to ensure equiluminance on the monitors used in the lab. Within a given participant, two colours were consistently used as the stimuli colours, while the third colour served as the experimental neutral colour. This neutral colour was also always used as the intervening task colour (see ‘*Task and Procedure*’ for details on the intervening task). Stimuli colours were randomly assigned on each trial, with the constraint that the two bar stimuli in one trial had a different colour and neither were the neutral colour. Which colour served as the neutral colour was counterbalanced across participants, so that the neutral colour changed from one participant to the next in the set order of orange - green - blue. After replacement of one participant, this resulted in orange as neutral colour for 8 participants, green for 9 participants, and blue for 8 participants.

For reasons explained in our Results section and visualised in **Figure 3**, we added an extra neutral colour in Experiment 2, so four colours were used: the original three from Experiment 1 (blue (rgb: 19, 146, 206), green (rgb: 101, 148, 14), and orange (rgb: 238,104, 60)), with magenta (rgb: 217, 103, 241) as the fourth colour. All four colours were calibrated prior to the experiment to ensure equiluminance on the monitors used in the lab. Within a given participant, two colours were consistently used as the stimuli colours, while the third and fourth colours served as the experimental neutral colours. One neutral colour was tied to the intervening task, just as in Experiment 1, while the other neutral was relevant for neither the intervening task, nor the working-memory task. Stimuli colours were randomly assigned on each trial, with the constraint that the two bar stimuli in one trial had a different colour and neither were either of the two neutral colours. Which of the four possible colours served as the neutral colour that was used for the intervening task was counterbalanced across participants, so that this neutral colour changed from one participant to the next in the set order of magenta - orange - green - blue. After replacement of three participants, this resulted in magenta as neutral colour for 7 participants, orange for 6 participants, blue for 6 participants, and green for 6 participants. Which colours served as the two stimuli colours and additional (task irrelevant) neutral colour were randomly drawn from the three remaining colours.

### Comparison to comparable conditions in prior unpublished experiments

In both Experiment 1 and Experiment 2, participants were required to evaluate a colour change. To assess the impact of this intervening task on our main outcomes we decided to combine the data from Experiments 1 and 2 (n=50, 512 trials per participant; 25.600 trials total) and compare them to two previously conducted experiments that had remained unpublished. In these experiments, Experiments 3 and 4 (n=50, respectively, 192 trials or 384 trials per participant; 14.400 trials total), participants performed an identical working-memory task with the same stimuli, but this time were not required to perform any additional intervening task based on the colour change. This comparison afforded us a high-powered between-subjects evaluation that directly informed us of the effect of having to engage with an unpredictive external stimulus (as shown in **Fig. 4**).

Experiments 3 and 4 also had an a-priori sample size of twenty-five healthy human volunteers, again recruited from the Vrije Universiteit Amsterdam. Experiment 3 initially included twenty-seven participants, two of whom were excluded for misunderstanding the task, resulting in the target sample size (age range: 18-31; 19 women, 5 men, 1 non-binary; 24 right-handed). Experiment 4 included twenty-five participants (age range: 18-28; 21 women, 4 men; 21 right-handed, 1 ambidextrous).

Half of the blocks in Experiments 3 and 4 were identical to Experiments 1 and 2, with one key difference that motivated this comparison: Experiments 3 and 4 each also included the same unpredictive colour change of the central fixation marker, but unlike in Experiments 1 and 2, participants were not required to do any intervening task based on this colour change, so the colour change could theoretically be disregarded. Consequently, Experiments 3 and 4 also did not include a neutral condition (the same experimental colours were used (blue, green, orange and magenta) and two were randomly assigned to the stimuli each trial). For transparency, we note that Experiments 3 and 4 also included conditions not present in Experiments 1 and 2. As these additional conditions did not allow for a meaningful comparison with Experiments 1 and 2, they were not included in the analyses here. The relevant (comparable) conditions involved 8 blocks of 24 trials in Experiment 3 and 8 blocks of 48 trials in Experiment 4. These experiments offered a valuable reference for assessing the impact of the intervening task on our main outcomes.

### Eye-tracking acquisition and preprocessing

In all four experiments, horizontal and vertical gaze positions of the participant’s right eye were continuously sampled using an EyeLink 1000 Plus at a rate of 1000 Hz throughout the experiment. The eye-tracker was positioned approximately 10 cm in front of the monitor, at approximately 60 cm away from the eyes. Prior to recording, the eye tracker was calibrated and validated using the built-in HV9 calibration module provided by the Eyelink software. The eye tracker was always re-calibrated exactly halfway through the experiment (after 8 blocks), and could additionally be re-calibrated in each break between consecutive blocks, if the signal was deemed of too poor quality.

Following acquisition, eye-tracker datafiles were converted from their original Eyelink Data Format (.edf) to an ASCII text file (.asc), and analysed in MATLAB R2024b using a combination of the Fieldtrip analysis toolbox^58^ and custom code. Custom code was used to detect blinks and remove them from the signal for 100 ms before and after each blink (by replacing these data with NaNs). After blink removal, data were epoched relative to onset of the unpredictive colour change of the central fixation marker.

### Saccade detection and visualisation

To detect saccades, a previously established and validated velocity-based method was employed^36^ (building on ^59^). In this approach, saccades are detected by finding those samples where the 1-dimensional gaze velocity exceeds a certain trial-based threshold. Gaze velocity in the horizontal (left/right) axis was obtained as the derivative of horizontal gaze position (i.e. the difference between the X-coordinates of consecutive samples) that was smoothed in the temporal dimension with a Gaussian-weighted moving mean filter with a 7-ms sliding window (using the built-in MATLAB function ‘smoothdata’). The velocity threshold was set at 5 times the median velocity in any given trial, which is consistent with earlier work^37,39,60,61^. A minimum delay of 100 ms between successive saccades was imposed to avoid counting the same saccade multiple times.

Saccade size (in degrees) and saccade direction (left/right) were calculated by estimating the difference between the pre-saccade gaze position (−50 to 0 ms before threshold crossing) and the post-saccade gaze position (50 to 100 ms after threshold crossing). Depending on the left/right direction of the saccade and the original left/right location of the colour-matching memory item, saccades were classified as ‘toward’ or ‘away’. After detecting and classifying the saccades, the time courses of saccade rates (in Hz) were determined by using a sliding time window of 100 ms, advancing in steps of 1 ms.

To also investigate the size of the saccades that contributed to our findings without imposing an arbitrary threshold, we additionally decomposed saccade rates into a time-size representation, showing the time courses of saccade rates as a function of the saccade size (as also done in ^36,60,62^). For this, we used successive saccade-size bins of 0.5 degrees visual angle in steps of 0.1 degree.

### Statistical analysis

For all experiments, we computed decision-time z-scores per participant (with decision times being defined as the time between dial onset and response initiation) and removed all trials with an absolute z-score greater than 3 from the performance analysis. For all analyses of Experiments 1 and 2 (performance and saccade data) we collapsed across the two different block types.

For the analysis of the performance data of Experiments 1 and 2, we employed linear mixed-effects modelling to examine differences in average performance between trials in which the external stimulus matched the eventually tested item, the untested item, or neither item (neutral condition). Linear mixed-effects models allow for the inclusion of participant-level variability in performance, by including random intercepts for participants. Two separate models were fitted, one for decision times (time in ms between dial onset and response initiation) and one for reproduction errors (the absolute angular difference between the reported and memorised item orientation). In both models, condition (three levels: match tested, match untested, match neutral) was included as a fixed effect, and random intercepts were specified for participants. Models were fitted using the built-in MATLAB function ‘fitlme’ with the following formulas:

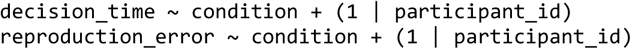

Post-hoc analyses of the performance data of Experiments 1 and 2 were conducted using paired-sample Student’s t-tests to examine differences in average decision times and reproduction errors for all three pairwise contrasts (tested vs. neutral, untested vs. neutral, tested vs. untested). For Experiment 2, comparisons with the neutral conditions were done on both neutral colour conditions combined. The resulting p-values were Bonferroni corrected for multiple comparisons. Effect sizes for each comparison were quantified using Cohen’s d. To quantify evidence for the absence of an effect in cases of non-significant results, Bayesian paired-sample t-tests were performed in JASP using default priors (Cauchy distribution centred at 0 with scale 0.707).

The performance data of Experiments 3 and 4 were analysed using paired-sample Student’s t-tests to examine differences in average decision times and reproduction errors between the match-tested and match-untested conditions. As only one contrast could be evaluated for each measure, the resulting p-values were not Bonferroni corrected for multiple comparisons. Effect sizes for each comparison were again quantified using Cohen’s d. To quantify evidence for the absence of an effect in cases of non-significant results, Bayesian paired-sample t-tests were performed in JASP using default priors (Cauchy distribution centred at 0 with scale 0.707).

For the between-subjects comparison of Experiments 1-2 with Experiments 3-4 (**Fig. 4C**), independent-sample Student’s t-tests were employed to compare the effect of condition on average decision times and reproduction errors. Effect sizes for each comparison were again quantified using Cohen’s d.

To statistically evaluate the time series saccade data, we employed a non-parametric cluster-based permutation approach^63^. The permutation analysis was conducted on the time period from 0 to 1500 ms after onset of the colour change, using Fieldtrip with default clustering settings (grouping adjacent time points that were significant in a mass-univariate comparison and summing their t-values to arrive at a cluster size). A permutation distribution of the largest cluster size was acquired by randomly permuting the condition labels of each participant’s trial-averaged time course data (i.e. randomly flipping the sign of the difference in rate of toward vs. away saccades) 10,000 times and identifying the size of the largest clusters observed in these randomised data after each permutation. The p-values of the clusters observed in the original data were calculated as the proportion of random permutations where the largest cluster was equal to or larger than the cluster that we observed in the original data.

Throughout this entire article, the following significance levels were used: *: p< 0.05, **: p < 0.01, ***: p < 0.001, ****: p < 0.0001 (this is also indicated in the figure captions).

## Acknowledgements

This research was supported by an NWO Vidi Grant by the Dutch Research Council (grant number 14721), and an ERC Starting Grant from the European Research Council (MEMTICIPATION, 850636) to F.v.E. In addition, we thank Holly Zhang and Fathima Shamsuddin for their help with the data collection of Experiments 3 and 4.

## Data, materials and software availability

All study data and analysis code will be made public prior to publication. The code for the experimental tasks is already publicly available on GitHub ([Experiment 1](https://github.com/annavanharmelen/Action-coupled-null-cue-experiment/tree/main), [Experiment 2](https://github.com/annavanharmelen/Action-coupled-null-cue-experiment/tree/extra-neutral), [Experiment 3](https://github.com/annavanharmelen/Location-by-colour-null-cue-experiment) and [Experiment 4](https://github.com/annavanharmelen/Location-by-colour-null-cue-experiment/tree/remove-location-cue)).

